# Interplay between *MYZUS PERSICAE-INDUCED LIPASE 1* and OPDA signaling in controlling green peach aphid infestation on *Arabidopsis thaliana*

**DOI:** 10.1101/2022.07.08.499389

**Authors:** Lani Archer, Hossain A. Mondal, Sumita Behera, Moon Twayana, Joe Louis, Vamsi J. Nalam, Jantana Keereetaweep, Zulkarnain Chowdhury, Jyoti Shah

## Abstract

*MYZUS PERSICAE-INDUCED LIPASE1* (*MPL1*) encodes a lipase in *Arabidopsis thaliana* that is required for controlling infestation by the green peach aphid (GPA; *Myzus persicae*), an important phloem sap-consuming insect pest. Previously, we demonstrated that *MPL1* expression was upregulated in response to GPA infestation, and GPA fecundity was higher on the *mpl1* mutant, compared to the wild-type (WT), and lower on *35S*:*MPL1* plants that constitutively expressed *MPL1* from the *35S* promoter. Here, we show that the *MPL1* promoter is active in the phloem and expression of the *MPL1* coding sequence from the phloem-specific *SUC2* promoter is sufficient to restore resistance to the GPA in the *mpl1* mutant. The GPA infestation-associated upregulation of *MPL1* requires *CYCLOPHILIN 20-3* (*CYP20-3*), which encodes a 12-oxo-phytodienoic acid (OPDA)-binding protein that is involved in OPDA signaling and is required for controlling GPA infestation. OPDA promotes *MPL1* expression to limit GPA fecundity, a process that requires *CYP20-3* function. These results along with our observation that constitutive expression of *MPL1* from the *35S* promoter restores resistance to the GPA in the *cyp20-3* mutant, and *MPL1* feedbacks to limit OPDA levels in GPA-infested plants, suggest that an interplay between *MPL1*, OPDA, and *CYP20-3* contributes to resistance to the GPA.

**Highlight:** Interaction between *MYZUS PERSICAE-INDUCED LIPASE 1* function in the phloem, and 12-oxo-phytodienoic acid (OPDA) and *CYCLOPHILIN 20-3*, which encodes an OPDA-binding protein that is involved in OPDA signaling, is involved in controlling green peach aphid infestation on *Arabidopsis thaliana*.

## Introduction

Aphids comprise a large group of hemipterans that include some of the most damaging pests of plants (Dixon, 1998; Blackman and Eastop, 2000; Dedryver *et al*., 2010). Aphids utilize their mouthparts, which are modified into stylets, to penetrate the sieve elements from which they consume phloem sap. Damage to plants results from the removal of phloem sap, changes in source-sink patterns that affect nutrient flow in the plant, and viral diseases vectored by aphids (Mittler and Sylvester, 1961; Kennedy *et al*., 1962; Matthews 1991; Dixon 1998; Girousse *et al*., 2005). The stylets take a largely intercellular path to the sieve elements. However, in their search for the sieve elements, they occasionally puncture non-vascular cells for gustatory cues (Pollard, 1973; Tjallingii and Esch, 1993). Thus, defenses against aphids could potentially be exerted at multiple levels, including at the phloem, the apoplastic space through which the stylets traverse, and the non-vascular cells penetrated by the stylets (Twayana *et al*., 2022).

Lipids and lipid metabolism influence plant interaction with aphids (Nalam *et al*., 2019; Twayana et al., 2022). In addition, as described below, aphid infestation impacts plant lipid metabolism. *Myzus persicae*, more commonly known as the green peach aphid (GPA), exhibits low reproduction and high mortality on tobacco (*Nicotiana tabacum*) that contain high levels of atypical sterols, which unlike cholesterol, the major sterol in aphids, lack double bonds in the sterol nucleus (Behmer *et al*., 2011). Also, GPA fecundity is lower on *Arabidopsis thaliana* mutants and transgenic lines that contain low levels of β-sitosterol than campesterol (Chen *et al*., 2020). Aphids lack the enzymatic machinery necessary for *de novo* synthesis of sterols, hence, presumably they can utilize plant-derived β-sitosterol but not atypical sterols to generate cholesterol (Behmer and Nes, 2003; Bouvaine *et al*., 2014). Host fatty acid desaturation also influences aphid infestation. GPA fecundity was lower on the Arabidopsis *ssi2* mutant, which is deficient in a stearoyl acyl-carrier protein (ACP)-desaturase that is involved in the synthesis of oleic acid (C18:1) from stearic acid (C18:0) in the plastids (Pegadaraju *et al*., 2005; Louis *et al*., 2010a; Li *et al*., 2021). Enhanced resistance to the GPA was also observed in the Arabidopsis *fad3* and *fad7* mutants that are deficient in endoplasmic reticulum- and plastid-localized ω-3 fatty acid desaturases, respectively (Li *et al*., 2021). These desaturases synthesize poly- unsaturated fatty acids. Likewise, loss of *FAD7* function in tomato (*Solanum lycopersicum*) enhanced resistance to the potato aphid (*Macrosiphum euphorbiae*) (Avila *et al*., 2012). The physiological basis of the impact of altered unsaturated fatty acid composition on plant-aphid interaction is unclear.

Aphid infestation impacts oxidized lipid (oxylipin) metabolism in plants. Levels of 9- hydroxyoctadecadienoic acid (9-HOD) and 9-hydroxyoctadecatrienoic acid (9-HOT), which are synthesized as a result of oxidation of poly-unsaturated fatty acids by 9-lipoxygenases (9-LOXs), increase in the phloem sap of Arabidopsis in response to GPA infestation (Nalam *et al*., 2012).

The increase in oxylipin levels was associated with the upregulation of *LOX5* expression in roots of GPA-infested plants (Nalam *et al*., 2012). GPA fecundity and feeding from sieve elements was reduced on the *lox5* mutant. Irrigation with 9-HOD enhanced GPA performance on the *lox5* mutant. Furthermore, 9-HOD when included in a synthetic diet increased GPA fecundity (Nalam *et al*., 2012). These oxidized lipids can be recovered from GPA reared on Arabidopsis (Nalam *et al*., 2013) and faba beans (*Vicia faba*) (Harmel *et al*., 2007). Since extracts from the GPA lack the ability to synthesize these oxidized lipids, it has been suggested that the GPA obtains these oxidized lipids from its diet (Harmel *et al*., 2007). These plant 9-LOX-derived oxylipins could function to stimulate aphid feeding and/or aphid reproduction. Alternatively, they may be involved in suppressing host defenses. α-dioxygenases encode another class of fatty acid oxidizing enzymes that influence aphid performance. In tomato and Arabidopsis, loss of α- dioxygenase-1 activity resulted in improved performance of the potato aphid and green peach aphid, respectively (Avila *et al*., 2013), thus suggesting that α-dioxygenase-synthesized oxylipins, or products thereof, function to limit aphid infestation.

Jasmonates are a group of cyclopentanone oxylipins that have important signaling functions in plant response to herbivores, including aphids (Howe and Jander, 2008; Erb *et al*., 2012; Louis and Shah, 2013). Jasmonic acid (JA) (Supplementary Fig. 1A and S1B) is one of the best studied cyclopentanone oxylipin with a role in plant-aphid interaction. Resistance against the potato aphid in tomato was enhanced by JA application (Cooper *et al*., 2005), and resistance in sorghum (*Sorghum bicolor*) against the greenbug aphid (*Schizaphis graminum*) and in Arabidopsis against the GPA were enhanced by methyl-JA (MeJA) application (Ellis *et al*., 2002; Zhu-Salzman *et al*., 2004). Furthermore, in comparison to the wild-type (WT), the Arabidopsis *cev1* (*constitutive expression of VSP1*) mutant, which contains elevated JA levels, was more resistant to the GPA (Ellis *et al*., 2002). Accordingly, the GPA performed better on the Arabidopsis *coi1* (*coronatine insensitive1*) mutant, which is deficient in JA signaling (Ellis *et al*., 2002; Mewis *et al*., 2005, 2006). Resistance in Arabidopsis against the cabbage aphid (*Brevicoryne brassicae*), and in barrel medic (*Medicago truncatula*) against the blue green aphid (*Acyrthosiphon kondoi*) conferred by *AKR* (*Acyrthosiphon kondoi resistance*), also invoke JA signaling (Gao *et al*., 2007; Kusnierczyk *et al*., 2011). In comparison, JA has a dichotomous contribution in the interaction of sorghum with sugarcane aphid (*Melanaphis sacchari*) (Grover *et al*., 2022). JA level was found to transiently increase in response to sugarcane aphid infestation in the resistant sorghum variety SC265. Sugarcane aphid feeding was restricted and the population curtailed on the JA-deficient mutant compared to the WT. Also, exogenous application of JA promoted sugarcane aphid feeding and colonization on SC265, thus suggesting that JA facilitates sugarcane aphid infestation on the sorghum variety SC265. However, sugarcane aphid settling was enhanced on the JA-deficient mutant (Grover *et al*., 2022).

12-oxo-phytodienoic acid (OPDA) (Supplementary Fig. S1A and S1B), which is an intermediate in the biosynthesis of JA, is also involved in plant defense signaling (Liu and Park, 2021). OPDA also contributes to defense against aphids. In the maize (*Zea mays*) inbred line Mp708, resistance to the corn leaf aphid (*Rhopalosiphum maidis*) was associated with the constitutively higher levels of OPDA (Varsani *et al*., 2019). Furthermore, OPDA application to a maize line that was deficient in the synthesis of JA, enhanced resistance to the corn leaf aphid, leading to the suggestion that OPDA functions independently of JA in promoting resistance to the corn leaf aphid. OPDA was found to promote defenses, for example the deposition of callose, in response to corn leaf aphid infestation (Varsani *et al*., 2019).

Considering that a substantial part of an aphid’s time on a plant is spent consuming phloem sap from the sieve elements, the phloem provides an important front for the exertion of defenses (Twayana *et al*., 2022). However, there is a lack of comprehensive knowledge on the molecular and physiological processes associated with phloem-based defense against aphids.

Previously, *MYZUS PERSICAE-INDUCED LIPASE 1* (*MPL1*), which encodes an α/β-fold lipase, was shown to be required for controlling GPA infestation on Arabidopsis (Louis *et al*., 2010b). *MPL1* expression was upregulated in response to GPA infestation and resistance against GPA was enhanced in *35S*:*MPL1* plants that constitutively expressed high levels of *MPL1* from the *Cauliflower mosaic virus 35S* gene promoter (Louis *et al*., 2010b). In comparison, GPA population was larger on *mpl1-1* and *mpl1-2*, which contained T-DNA insertions within the *MPL1* coding sequence. The *mpl1-1* allele also attenuated the *ssi2*-conferred enhanced resistance to the GPA (Louis *et al*., 2010b). Here, we demonstrate that GPA infestation results in the upregulation of *MPL1* promoter activity in the phloem and *MPL1* expression from the phloem-specific *SUCROSE-PROTON SYMPORTER 2* (*SUC2*) promoter (Stadler and Sauer, 2019) is sufficient to restore resistance to the GPA in the *mpl1-1* background. We provide physiological and genetic evidence that suggests an interplay between *MPL1* and OPDA signaling in defense against the GPA. While *MPL1* functions to limit OPDA accumulation, the OPDA-binding cyclophilin CYP20-3, which is required for OPDA signaling, controls upregulation of *MPL1* expression in response to GPA infestation and like *MPL1* is required for controlling GPA infestation.

## Materials and Methods

### Plant genotypes and cultivation

Arabidopsis was cultivated under a 14-h light (80 - 100 µE m^−2^ s^−1^) and 10-h dark regime at 22°C (Nalam *et al*., 2016). The *35S*:*MPL1*, *mpl1-1* (Salk_101919) and *mpl1-2* (Salk_082589) lines are in the accession Columbia-0 (Col-0) (Louis *et al*., 2010b). The *MPL1*p:*UidA* plants (Alam *et al*., 2022) express the GUS-encoding *Escherichia coli UidA* from a 1785 bp *MPL1* promoter. The *cyp20-3* (SALK_001615C) (Park *et al*., 2013b), *coi1-*17 (Suza and Staswick, 2008; Staswick, 2009), *jar1-1* (Staswick and Tiryaki, 2004), *pxa1* (Zolman *et al*., 2001), *acx1 acx5* (Schilmiller *et al*., 2007), and *lox2/3/4/6* quadruple mutant (Yang *et al*., 2020; Chauvin *et al*., 2013), which is referred in this study as *lox*(q), are in the Col-0 background. *Lox*(q) is derived from the *lox3* (Salk_062064), *lox4* (Salk_071732), *lox6* (Salk_083650) and a previously described non-sense mutation containing *lox2-1* mutant (Glauser *et al*., 2009). The *aos* mutant (stock # CS6149; https://abrc.osu.edu/) is in the Columbia *glabra1* background. In order to obtain seeds from the homozygous *lox*(q) and *aos* plants, which are male sterile due to JA-deficiency, the floral buds were treated daily with MeJA (50 µM) to promote pollen development. The *cyp20-3 35S:MPL1* plants were obtained by crossing a *cyp20-3* with a *35S*:*MPL1* plant. The resultant F1 plants were allowed to set seeds and the F2 progeny screened for the presence of the *35S*:*MPL1* transgene by PCR with the primer set 35S-F1 and MPL1-CDS-Stop-R1 and for plants homozygous for the *cyp20-3* allele by PCR with the primer set T-DNA-LB and CYP20-3-R1 (Supplementary Table S1).

### Generation of SUC2p:MPL1 plants

A pMDC85 derivative (Gottwald *et al*., 2000) in which the *35S* promoter was replaced by the phloem-specific *SUC2* promoter (Stadler and Sauer, 2019), was used to generate the *SUC2*p:*MPL1* chimera in which the *MPL1* coding sequence is expressed from the *SUC2* promoter. The LR clonase system (https://www.thermofisher.com) was used to mobilize the *MPL1* coding sequence contained in a pCR8 vector (Louis *et al*., 2010b) into the pMDC85- *SUC2*p vector. The resultant *SUC2*p:*MPL1* construct, in the pMDC85 background was electroporated into *Agrobacterium tumefaciens* GV3101, from which T-DNA containing the *SUC2*p:*MPL1* chimera was transformed into the *mpl1-1* mutant by the floral-dip method (Zhang *et al*., 2006). Hygromycin resistant seedlings were identified by germinating seeds on Murashige and Skoog agar plates containing 20 mg ml^-1^ of hygromycin. The presence of the chimeric construct in the hygromycin-resistant seedlings was confirmed by PCR with the primer set SUC2p-F1 and MPL1-CDS-Stop-R1 (Supplementary Table S1). Expression of *MPL1* from *SUC2*p:*MPL1* was confirmed by RT-PCR with the MPL1-F1 and MPL1-R1 primer set (Supplementary Table S1).

### OPDA treatment of Arabidopsis

Leaves were treated with 50 µM OPDA dissolved in 0.2% ethanol, and as control with 0.2% ethanol, by brushing 10 µL of solution on leaves. Leaf samples for RNA extraction were harvested 24 h later.

### Rearing the green peach aphid and aphid bioassays

The GPA was reared on a 50:50 mix of radish (Radish Early Scarlet Globe; Main Street Seed & Supply, catalog number 13307-13) and mustard plants (Florida Mustard Broad Leaf; Main Street Seed & Supply, catalog number 12501-13), as previously described (Nalam *et al*., 2018). For aphid fecundity assays, 20 adult aphids were placed on each Arabidopsis plant and the number of nymphs born over a 48h period determined. Each experiment included 10-15 plants of each genotype. Aphid fecundity was calculated as the number of nymphs born, per adult, per day. For experiments involving OPDA treatment, two adult aphids were placed on each leaf, 24 h after OPDA/ethanol treatment, following which each leaf was individually caged in a 2 ml microfuge tube with holes to allow gas exchange. The number of nymphs born were determined 48h later.

A synthetic diet (Mittler and Dadd, 1965) supplemented with different concentrations of OPDA and JA was used to study the impact of OPDA and JA on aphid reproduction (Louis *et al*., 2010a). Two adult aphids were allowed to feed on each feeding chamber with the synthetic diet contained between two layers of parafilm. The total population size (adults plus nymphs) was measured four days later. Four biological replicates were included for each treatment.

### RNA isolation and gene expression

Total RNA was extracted from leaves by grinding frozen leaf samples in an acid guanidinium thiocyanate/phenol/chloroform mix (Chomczynski and Sacchi, 1987). After treatment with RQ1 DNase (Promega, Madison, WI; www.promega.com) to degrade DNA, GoScript^TM^ reverse transcriptase (Promega, Madison, WI; www.promega.com) along with oligo-dT 18 mer primer (New England Biolabs, Ipswich, MA; www.neb.com) was used to synthesize cDNA. For monitoring *MPL1* and *CML46* expression, a minimum of three biological and two technical replicates were analyzed for each genotype/treatment. Each biological replicate contained RNA extracted from two leaves. An iTaq Universal SYBR Green mix (Bio-Rad Lab, Hercules, CA; www.bio-rad.com) was used for qPCR with an ECO Illumina Real-Time PCR system (Illumina, San Diego, CA; www.illumina.com). The presence of a single PCR product was confirmed by a melt curve. Gene expression levels were normalized by subtracting the cycle threshold (C_T_) value of the control elongation factor (EF) gene At1g07920 (Singh *et al*., 2012; Jaouannet *et al*., 2015) from that of the gene of interest. Fold changes in gene expression were calculated based on the ΔΔC_T_ method (Livak and Schmittgen, 2001). Gene-specific primers used for qPCR are listed in Supplementary Table S1.

### Histochemical staining

X-gluc (5-bromo-4-chloro-3-indolyl-b-D-glucopyranosiduronic acid) (Gold Biotechnology, St. Louis, MO; https://www.goldbio.com/) was used as the substrate for the histochemical analysis of GUS activity. Freshly harvested leaf tissues were placed in a 100 mM phosphate buffer solution (pH 7.0) containing 50 µM each of potassium ferrocyanide and potassium ferricyanide, 0.1% Triton-X-100, and X-gluc (1 mg ml^-1^). After vacuum infiltration to facilitate substrate penetration into the leaves, the samples were incubated at 37°C to allow for GUS to hydrolyze X-gluc leading to the production of blue precipitate of chloro-bromoindigo. After the staining, the tissues were washed with multiple changes of 70% ethanol to remove chlorophyll, the samples were visualized under a bright field microscope.

### Oxylipin analysis

For hydroxylipid analysis, leaf tissues were processed and lipids were extracted as previously described (Göbel *et al*., 2002; Nalam, 2012) and resuspended in 100 µl methanol:water (80:20 v/v). A Nucleosil 120-5 C18 column (4.6 x 150 mm, 5 μm Macherey–Nagel, Bethlehem, PA, USA) was used as part of Reverse phase-HPLC (Agilent 1100 HPLC coupled to a UV diode array detector) to purify the hydroxylipids. A binary gradient solvent system consisting of methanol:water:acetic acid (80:20:0.1 v/v/v; solvent A) and methanol:acetic acid (100:0.1 v/v; solvent B) was utilized at a flow rate of 0.18 ml/ min. The hydroxylipids were further resolved by normal-phase HPLC (Zorbax Rx-SIL column, 2.1 x 150 mm, 5 μm, Agilent, Waldbronn, Germany) a flow rate of 0.125 ml min^-1^ with a solvent system that contained hexane: isopropanol: trifluoroacetic acid (100:1:0.02 v/v/v). Chiral phase HPLC (OD-H column; 150 x 2.1 mm, 5 µm particle size; Diacel, Osaka, Japan) with a solvent mix consisting of n of n-hexane:2-propanol:acetic acid (100:12.5:0.05, v/v/v) at a flow rate of 0.1 ml min^-1^ was used to distinguish the *S* from *R* isomers The *S* isomer was the most abundant compared to the *R* isomer, thus confirming an enzymatic source for vast majority of the hydroxylipids.

OPDA and JA were extracted from Arabidopsis leaves, and then purified, derivatized and quantified against a D5-JA standard by gas chromatography-mass spectrometry (Kilaru *et al*., 2007; Park *et al*., 2013a).

### Statistical analysis

Student’s *t*-test (two-tail) was used to assess the significance of variance (*P* < 0.05) when comparing two genotypes or treatments. ANOVA following the General Linear Model was used when comparing multiple genotypes and/or treatments to each other. Tukey’s test was used to assess the significance of variance (*P* < 0.05).

## Results

### MPL1 functions in the phloem to control GPA infestation

Reverse transcription-quantitative PCR (RT-qPCR) was used to validate the previously reported upregulation of *MPL1* expression in response to GPA infestation (Louis *et al*., 2010b). As shown in Supplementary Fig. 2A, *MPL1* expression was significantly higher in leaves from GPA infested compared to uninfested WT Arabidopsis plants. A similar increase in *MPL1* expression was not observed in the previously described *mpl1-1* mutant (Louis *et al*., 2010b). In comparison, *MPL1* expression was constitutively elevated in leaves of the *35S*:*MPL1* plant in the *mpl1-1* background. As previously demonstrated (Louis *et al*., 2010b), GPA performed significantly better on the *mpl1-1* mutant than the WT plant (Supplementary Fig. S2B). GPA fecundity was 50% higher on *mpl1-1* than the WT. In comparison, GPA fecundity was 25% lower on *35S:MPL1* compared to the WT plants.

To study the spatial expression pattern of *MPL1* in response to GPA infestation, β- glucuronidase (GUS) activity was followed in *MPL1*p:*UidA* plants in which the GUS-encoding *Escherichia coli UidA* was expressed from the *MPL1* promoter (*MPL1*p). GUS activity was monitored using the synthetic substrate X-gluc before and after GPA infestation. As shown in Fig. 1A, in GPA uninfested plants, GUS activity, reflective of the *MPL1* promoter activity, was negligible to very weak in most parts of the leaf, except for the hydathodes. In contrast, in response to GPA infestation, GUS activity was higher in the vascular tissues compared to rest of the leaf (Fig. 1A and 1B). Within the vasculature, strongest expression was observed in the phloem (Fig. 1B).

**Fig. 1.**
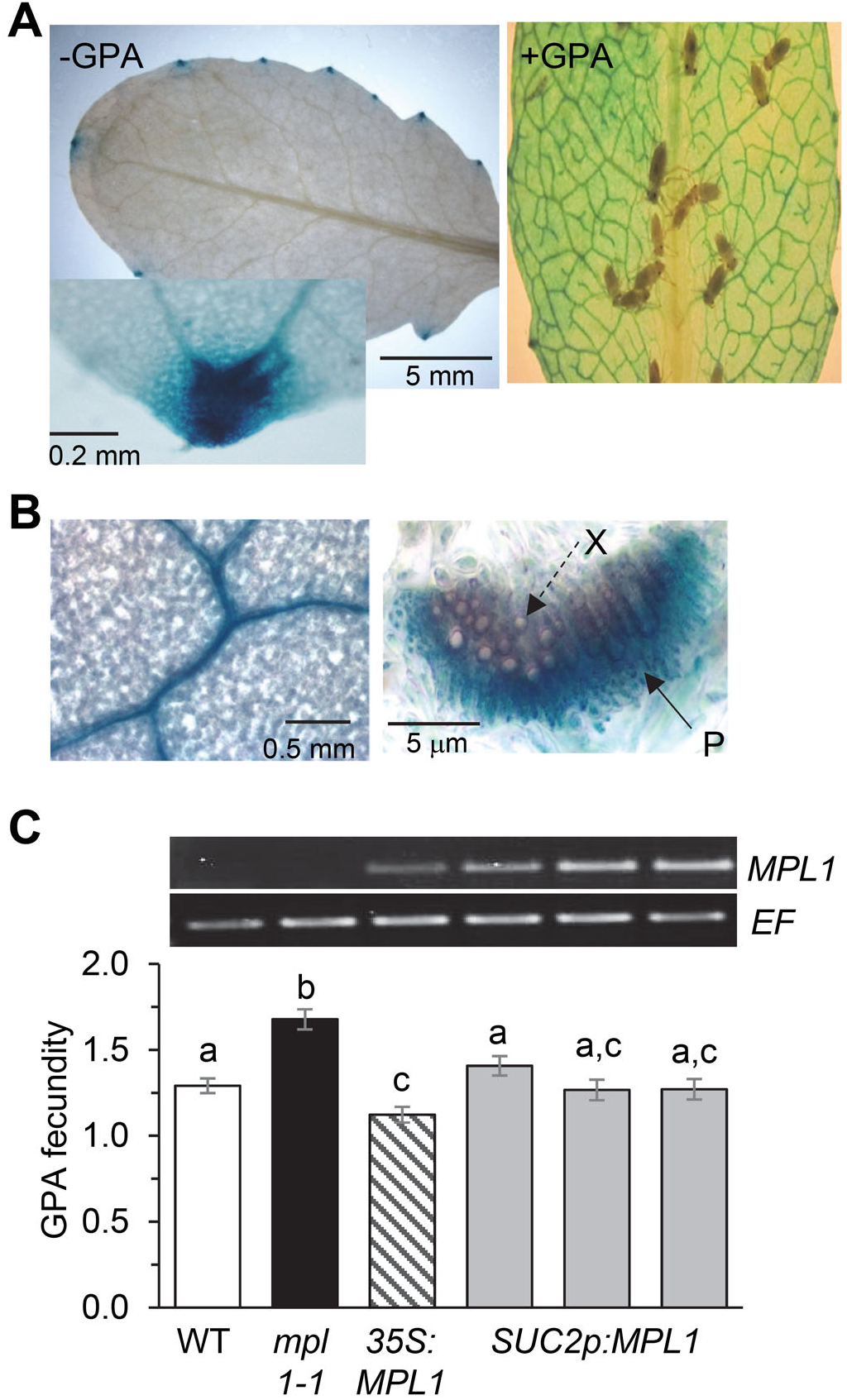
MPL1 functions in the phloem to limit green peach aphid infestation. **(A)** Histochemical analysis for GUS activity (blue color) in uninfested (left panel) and a GPA-infested leaf (right panel) of *MPL1*p:*UidA* in which expression of the GUS-encoding *UidA* is driven from the *MPL1* promoter. The inset in the left panel shows strong staining in the hydathodes of uninfested plants. **(B)** Left panel: Histochemical staining showing GUS activity in the veins of a GPA-infested leaf. Right panel: Cross section through the vein showing GUS activity in the phloem (P). X, xylem. **(C)** *Upper panel*: Constitutive *MPL1* expression in uninfested leaves of three independent *SUC2p*:*MPL1* lines in the *mpl1-1* background. The wild-type (WT), *mpl1-1* and *35S:MPL1* (in *mpl1-1* background) provided the control genotypes. *Lower panel*: GPA fecundity in the above plants. Error bars represent ± SE (n=10-12). Different letters above the bars represent values that are significantly different from each other (P<0.05; ANOVA).

To test if *MPL1* function in the phloem has a role in controlling GPA infestation, *mpl1-1 SUC2*p:*MPL1* plants were generated in which the *MPL1* coding sequence was expressed from the phloem-specific Arabidopsis *SUC2* promoter (Stadler and Sauer, 2019). GPA fecundity was evaluated on the *mpl1-1 SUC2*p:*MPL1* and as control on the *mpl1-1* and WT plants. Compared to *mpl1-1*, GPA fecundity was significantly lower on the *mpl1-1 SUC2*p:*MPL1* plants, and comparable to that on the WT plants (Fig. 1C), thus indicating that *MPL1* expression from the *SUC2* promoter is sufficient to restore WT-level of resistance in the *mpl1-1* background. Taken together, these results indicate that MPL1 function in the phloem has a critical role in defense against the GPA.

### MPL1 limits OPDA accumulation in response to GPA infestation

GPA infestation is accompanied by an increase in lipid oxidation (Gosset *et al*., 2009; Nalam *et al*., 2012). The levels of 13-LOX-derived 13-hydroperoxy and 13-hydroxy fatty acids (13-HP- FAs) are higher in GPA-infested compared to uninfested leaves (Fig. 2). 13-hydroperoxy linolenic acid feeds into the biosynthesis of JA and OPDA (Supplementary Fig. S1B), which have been implicated in plant defense and in promoting the accumulation of metabolites that are detrimental to aphids (Ellis *et al*., 2002; Mewis *et al*., 2005, 2006; Mikkelsen *et al*., 2003; Adio *et al*., 2011; Varsani *et al*., 2019; Grover *et al*., 2020). We therefore tested if GPA infestation increases levels of JA and OPDA in leaves of Arabidopsis. As shown in Fig. 3A, GPA infestation was not accompanied with significant changes in the content of JA in the WT plant.

**Fig. 2.**
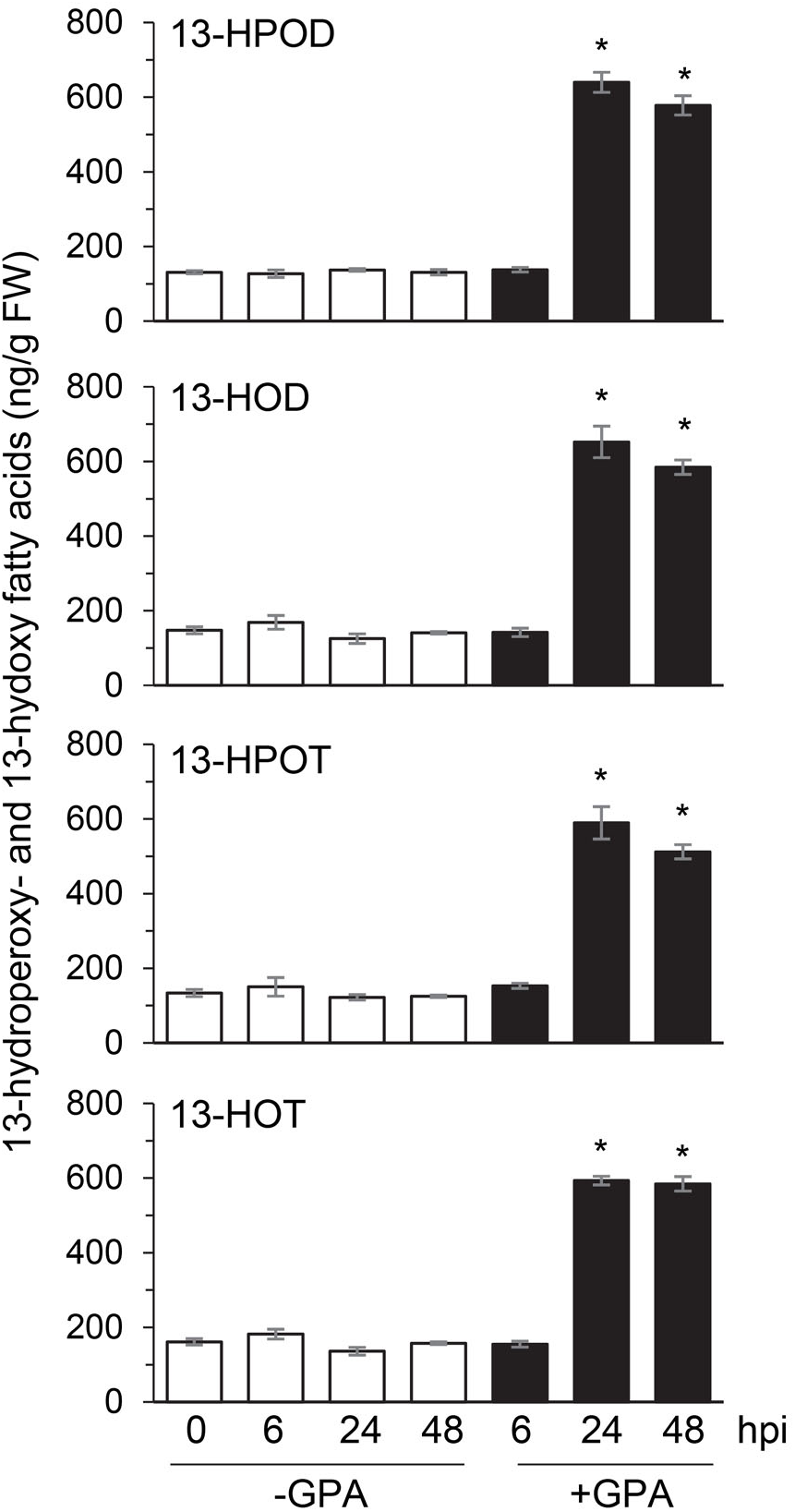
Changes in 13-LOX-derived oxidized lipids in Arabidopsis leaves in response to green peach aphid infestation. Levels of 13-hydroperoxy- and 13-hydoxy-fatty acids in leaves of green peach aphid-infested (+GPA) and uninfested (-GPA) wild-type accession Col-0 plants. Twenty adulty aphids were released on each plant. Leaf tissues were collected at the indicated hours post infestation (hpi), and as control from uninfested plants. Mean levels of 13-hydroperoxydienoic acid (13-HPOD), 13-hydroxydienoic acid (13-HOD), 13-hydroperoxytrienoic acid (13-HPOT), and 13-hydroxytrienoic acid (13-HOT) are presented as ng g^-1^ fresh weight (FW) of leaves. Error bars represent ± SE (n=5). An asterisk (*) indicates significant difference (P < 0.05; *t*-test) from the uninfested tissue for the indicated time.

**Fig. 3.**
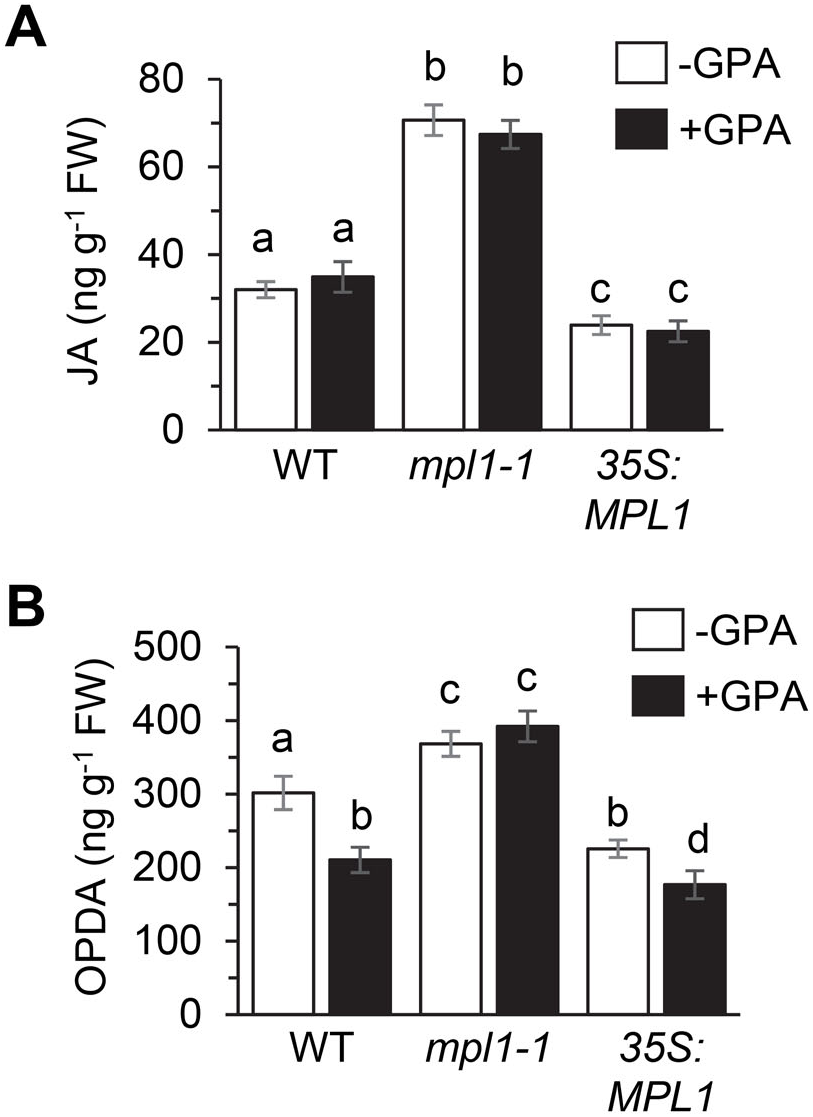
*MPL1* impacts 12-oxo-phytodienoic acid and jasmonic acid levels in Arabidopsis leaves. **(A)** Jasmonic acid (JA) and **(B)** 12-oxo-phytodienoic acid (OPDA) levels in leaves of wild-type (WT), *mpl1-1* and transgenic plants constitutively expressing *MPL1* from the *CaMV 35S* promoter. Leaves for JA and OPDA measurements were harvested from uninfested and GPA- infested plants at 24 hpi. Error bars represent ± SE (n=3). Different letters above bars represent values that are significantly different from each other (P<0.05; *t*-test).

However, contrary to expectations, OPDA content was significantly lower in the leaves of WT GPA-infested compared to the uninfested plants (Fig. 3B). In comparison to the WT plants, this reduction in OPDA content was not observed in the GPA-infested *mpl1-1* mutant (Fig. 3B), thus suggesting that *MPL1* negatively influences OPDA accumulation in response to GPA infestation. In concurrence with a recent report (Alam *et al*., 2022), basal content of OPDA and JA were also higher in the GPA-uninfested *mpl1-1* and lower in *35S:MPL1* compared to the WT leaves, thus implicating the involvement of *MPL1* in limiting OPDA and JA accumulation in Arabidopsis.

### The OPDA-binding CYP20-3 is required for controlling GPA infestation

Plant-derived oxylipins have been recovered from the gut of the GPA (Harmel *et al*., 2007; Nalam *et al*., 2013). In addition, OPDA which is a strong electrophile (Mueller and Berger, 2009), is toxic to lepidopteran larvae (Vollenweider *et al*., 2000). We therefore tested if the inclusion of OPDA, and as a control JA, in a synthetic diet affected GPA population. As shown in Fig. 4, inclusion of either OPDA or JA did not affect GPA population on a synthetic diet. We next tested if OPDA or JA accumulation and signaling influence Arabidopsis-GPA interaction. Aphid fecundity was compared on mutants previously shown to be deficient in steps leading to the synthesis of OPDA and JA (Supplementary Fig. S1B), and mutants defective in OPDA and JA signaling. As shown in Fig. 5A, compared to the WT, GPA fecundity was significantly higher on the *lox*(q) mutant, which is deficient in the four 13-LOXs encoded by *LOX2*, *LOX3*, *LOX4* and *LOX6* that catalyze the first oxidation step in the synthesis of OPDA and JA, and the *aos* (*allene oxide synthase*) mutant, which is defective in the synthesis of allene oxide, a precursor of OPDA and JA, thus suggesting that accumulation of OPDA/JA, or products thereof, is essential for controlling GPA infestation.

**Fig. 4.**
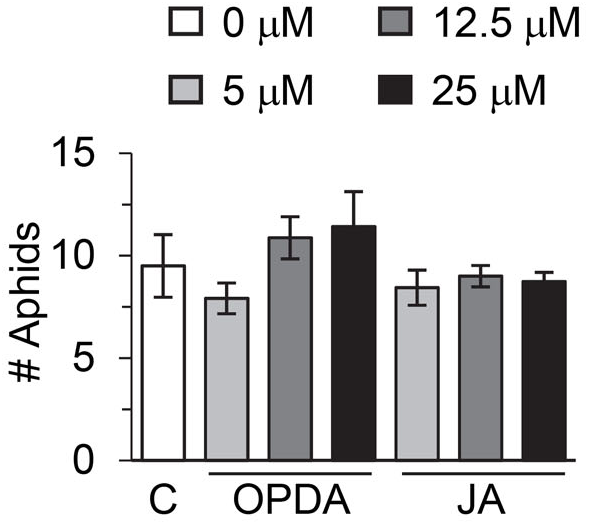
Green peach aphid performance on synthetic diet supplemented with OPDA and JA. Insect numbers (adult plus nymphs) were determined 4 days post release of two adults on a synthetic diet supplemented with different concentrations of OPDA and JA. Diet lacking any supplements provided the control (C). Error bars represent ± SE (n=4). Compared to the control (C), none of the treatments had a significant impact on GPA numbers (P>0.05; *t*-test).

**Fig. 5.**
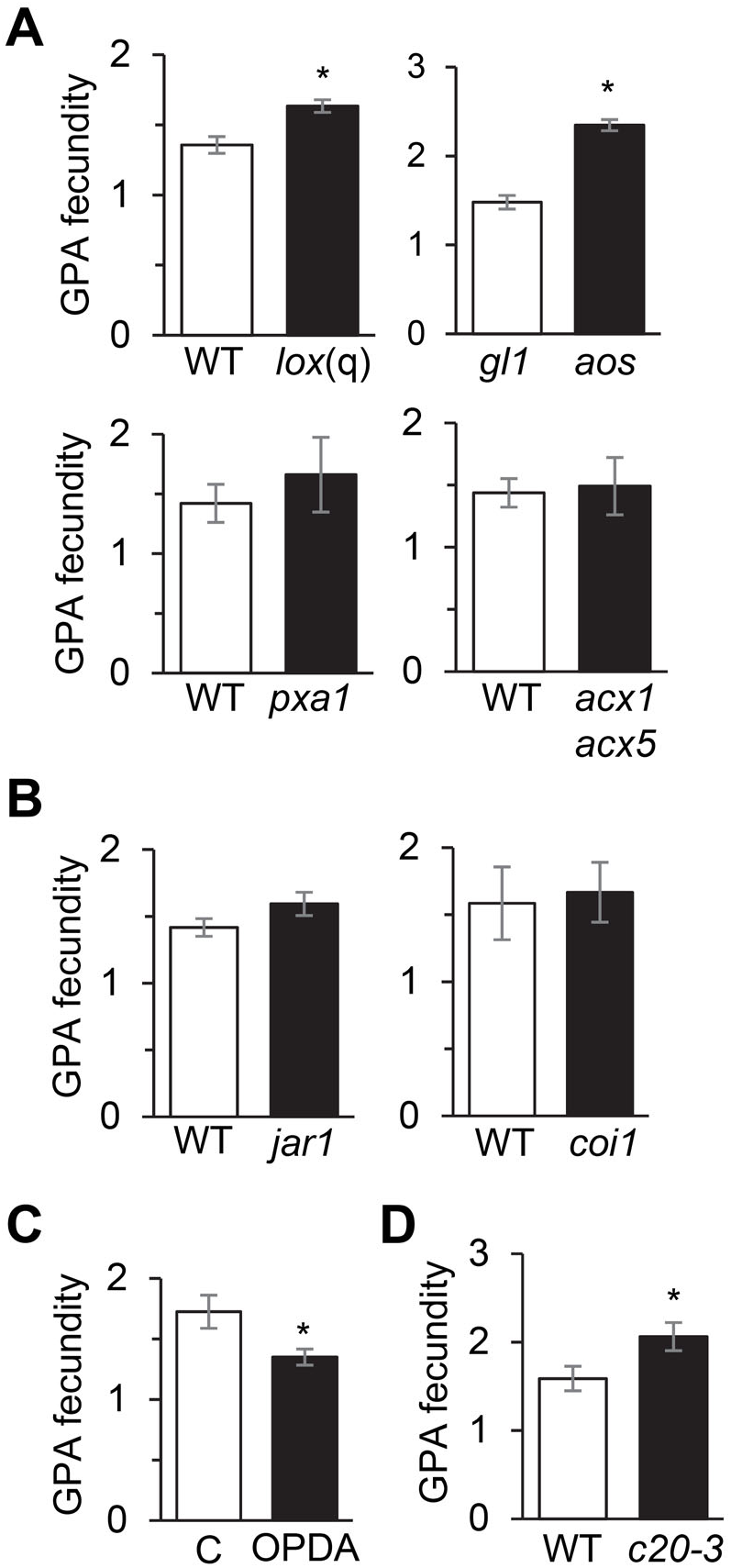
*CYP20-3* contributes to Arabidopsis defense against the green peach aphid. **(A)** GPA fecundity on mutants defective in OPDA and JA biosynthesis. The *lox*(q), *pxa1*, and *acx1 acx5* mutants are in the wild-type (WT) accession Columbia (Col-0), and the *aos* mutant is in the Columbia *glabra1* (*gl1*) background. Error bars represent ± SE (n=10). Asterisks (*) represent values that are significantly different between genotypes (P<0.05; *t*-test). **(B)** GPA fecundity on the JA-Ile deficient *jar1* and the JA signaling deficient *coi1* mutant. WT Col-0 provided the control genotype for *jar1* and *coi1*. Error bars represent ± SE (n=10). None of the values were significantly different from the corresponding control genotype (P>0.05; *t*-test). **(C)** GPA fecundity on OPDA (50 µM)-treated leaves of WT plant. Leaves treated with 0.2% ethanol, which was used to solubilize OPDA, provided the control (C). Error bars represent ± SE (n=10). An asterisk (*) indicates a value that is significantly different from the control treatments (P<0.05; *t*-test). **(D)** GPA fecundity on the OPDA signaling deficient *cyp20-3* (*c20-3*) mutant and WT Col-0. Error bars represent ± SE (n=15). An asterisk (*) indicates a value that is significantly different from the WT (P<0.05; *t*-test).

In contrast to the higher GPA fecundity on the *lox*(q) and *aos* mutants, GPA fecundity on the *pxa1* (*peroxisomal ABC transporter1*) mutant (Fig. 5A), which is deficient in JA biosynthesis due to impaired transport of OPDA into peroxisomes (Theodoulou *et al*., 2005), was comparable to that on the WT plants. Similarly, GPA fecundity was comparable on the WT and the *acx1 acx5* double mutant (Fig. 5A), which lacks acyl-CoA oxidase activities that catalyze the terminal oxidation steps in JA synthesis (Schilmiller *et al*., 2007). Also, compared to the WT plants, GPA fecundity was not significantly different between the WT and the JA-Ile-deficient *jar1* mutant (Fig. 5B) (Staswick and Tiryaki, 2004), and the WT and the *coi1* mutant (Fig. 5B), which is defective in the JA-Ile coreceptor (Katsir *et al*., 2008). Thus, we conclude that JA accumulation and signaling are not critical to Arabidopsis defense against the GPA. However, we cannot rule out that JA functions redundantly with other mechanisms in Arabidopsis defense against the GPA and might have a contribution under other growth/environment conditions.

OPDA, which is synthesized in the chloroplast, although an intermediate in JA synthesis, is also a signaling metabolite (Taki *et al*., 2005; Mueller and Berger, 2009). OPDA promotes resistance to corn leaf aphid, brown planthopper (*Nilaparvata lugens*), and Hessian fly (*Mayetiola destructor*) in maize, rice and wheat, respectively (Guo *et al*., 2014; Cheng *et al*., 2018; Varanasi *et al*., 2019; Grover *et al*. 2020). OPDA application also promoted resistance to GPA in Arabidopsis (Fig. 5C). In Arabidopsis, the chloroplast-localized OPDA binding protein CYP20-3 is required for OPDA signaling (Park *et al*., 2013b, Cheong *et al*., 2017). Considering that GPA infestation impacts OPDA accumulation and *MPL1* influences OPDA levels, we tested if *CYP20-3* influences the interaction between Arabidopsis and the GPA. As shown in Fig. 5D, GPA fecundity was higher on *cyp20-3* compared to the WT, thus leading us to conclude that *CYP20-3*, and hence OPDA signaling, have important functions in limiting GPA infestation on Arabidopsis.

### CYP20-3 promotes MPL1 expression in response to GPA infestation

To test the relationship between *CYP20-3* and *MPL1*, we analyzed the impact of *cyp20-3* on *MPL1* expression in response to GPA infestation. Expression of the OPDA-inducible *CML46* (At5g39670) (Taki *et al*., 2005), which encodes an EF hand Ca^2+^-binding protein, was monitored as a positive control in these experiments. As shown in Fig. 6A, GPA infestation resulted in the upregulation of *CML46* expression. This induction of *CML46* expression required *CYP20-3*, thus confirming the activation of OPDA signaling in GPA-infested plants. Similar to *CML46*, the upregulation of *MPL1* expression in response to GPA infestation was attenuated in *cyp20-3* relative to the WT, thus indicating that signaling through *CYP20-3* promotes *MPL1* expression in response to GPA infestation. Indeed, like *CML46*, OPDA application stimulated *MPL1* expression in the WT, but not in the *cyp20-3* mutant plants (Fig. 6B), thus further supporting a role for *CYP20-3* and hence OPDA signaling in promoting *MPL1* expression. The absence of *MPL1* induction in GPA-infested *lox*(q) mutant, further supports a role for a 13-LOX dependent product, presumably OPDA, in promoting *MPL1* expression in response to GPA infestation (Supplementary Fig. S3).

**Fig. 6.**
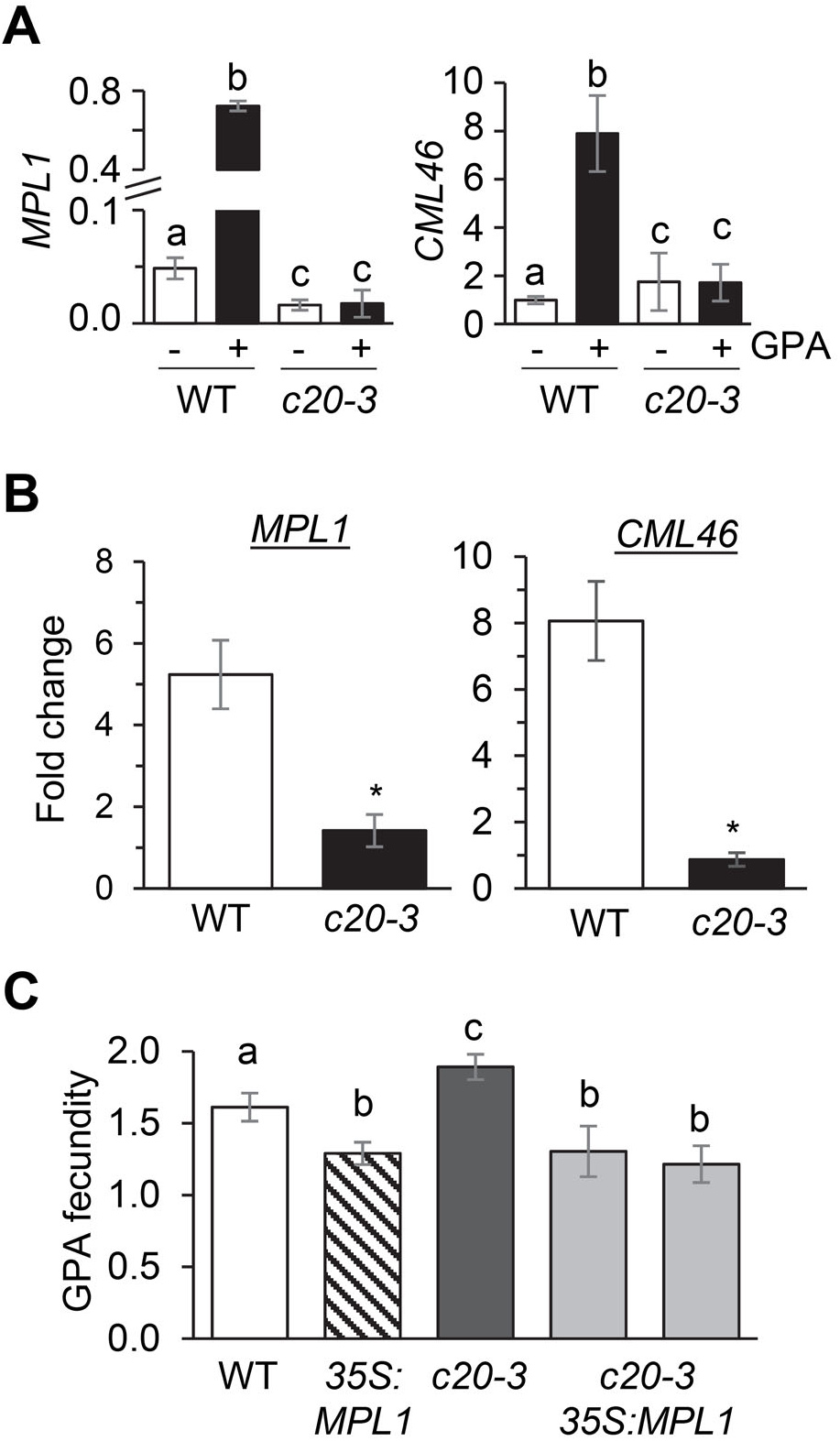
*CYP20-3* promotes *MPL1* expression to curtail GPA infestation. **(A)** *MPL1* and *CML46* levels (x 10^-2^) relative to the elongation factor gene At1g07920 at 24 hpi with the GPA (+GPA) and as control in uninfested (-GPA) WT and *cyp20-3* (*c20-3*) plants. Error bars represent ± SE (n=3). Different letters above bars represent values that are significantly different from each other (P<0.05; ANOVA). **(B)** Fold increase in *MPL1* and *CML46* expression in OPDA-treated relative to 0.2% ethanol-treated WT and *c20-3* leaves at 24 hpt. Error bars represent ± SE (n=3). Asterisks (*) indicate values that are significantly different from the WT (P<0.05; *t*-test). **(C)** GPA fecundity on the WT, *35S:MPL1*, *c20-3,* and *cyp20-3 35S:MPL1* (*c20-3 35S:MPL1*) plants. Error bars represent ± SE (n=10). Different letters above bars represent values that are significantly different from each other (P<0.05; ANOVA).

### Constitutive expression of MPL1 overcomes the cyp20-3 defect to limit GPA fecundity

To further characterize the relationship between *MPL1* and *CYP20-3*, *cyp20-3* was crossed with *35S*:*MPL1* to generate *cyp20-3 35S*:*MPL1* plants. Aphid fecundity was tested on the *cyp20-3 35S*:*MPL1* and as control the *cyp20-3*, *35S*:*MPL1* and WT plants. We hypothesized that if *MPL1* functions downstream of *CYP20-3* then the constitutive expression of *MPL1* from the *35S* promoter should overcome the *cyp20-3* defect to limit GPA fecundity in the *cyp20-3 35S*:*MPL1* plants. However, if *MPL1* functioned upstream of *CYP20-3* participation in defense against the GPA then *35S*:*MPL1* would be unable to overcome the *cyp20-3* defect to limit GPA fecundity in the *cyp20-3 35S*:*MPL1* plants. As shown in Fig. 6C, in comparison to the *cyp20-3* mutant, GPA fecundity on the *cyp20-3 35S*:*MPL1* plants was lower and comparable to that on the *35S*:*MPL1* plants, thus indicating that *35S*:*MPL1* overcomes the need of *CYP20-3* in limiting GPA infestation.

## Discussion

While studying the function of *MPL1* in Arabidopsis defense, we uncovered MPL1’s involvement in limiting levels of OPDA and JA. OPDA and JA levels were higher in *mpl1* mutant and lower in *35S:MPL1* plants that constitutively express *MPL1*. How MPL1, which is a lipase, limits OPDA and JA levels is unclear. Considering that MPL1 has homology to lipases involved in lipid droplet biogenesis/metabolism (Chapman *et al*., 2012) and loss of *MPL1* function affects lipid droplet number in *mpl1* (McLinchie, 2017), it is plausible that MPL1 limits the availability of lipids for oxylipin synthesis. It is equally plausible that a MPL1-dependent process promotes the metabolism of OPDA into a biologically more active derivative.

Alternatively, a MPL1-dependent mechanism could promote OPDA turnover. Hormone turnover can have important role in signaling as has been shown for JA-Ile. In plants with reduced JA-Ile turnover due to mutations that affect jasmonoyl-*L*-isoleucine-12-hydroxylase activity, while JA-Ile levels were higher (Kitaoka *et al*., 2011), these plants lacked wound-induced growth inhibition and exhibited enhanced susceptibility to insects, both phenotypes that are reminiscent of JA-Ile deficiency (Poudel *et al*., 2016). These plants expressed the JA signaling repressing *JAZ* genes at elevated levels, thus indicating a block in JA signaling resulting from reduced turnover of JA-Ile.

In higher plants, JA biosynthesis occurs in three different compartments. As indicated in Supplementary Fig. S1B, the early steps are initiated in the plastid where α-linolenic acid is converted to OPDA by the sequential action of 13-lipoxygenase (13-LOX), allene oxide synthase, and allene oxide cyclase (Wasternack and Song, 2017). OPDA is further converted to 3LJoxoLJ2 (2′(Z)LJpentenyl)LJcyclopentaneLJ1LJoctanoic acid (OPC-8:0) and finally to JA after three cycles of β-oxidation in the peroxisomes (Li *et al*., 2005; Fig. 7). *LOX*, *AOS* and *AOC*, which are involved in the early steps leading to biosynthesis of OPDA and JA are expressed in the vasculature (Kubigsteltig *et al*., 1999; Vellosillo *et al*., 2007; Chauvin *et al*., 2013). In addition, the LOX, AOS and AOC proteins are present in the phloem (Hause *et al*., 2003; Stenzel *et al*., 2003), OPDA and JA accumulate in cells associated with the vasculature (Mielke *et al*., 2011; Stenzel *et al*., 2003), and OPDA is a phloem-translocated metabolite (Schulze *et al*., 2019). Whether the impact of MPL1 on limiting OPDA/JA accumulation is exerted in the phloem *per se* is not known. However, considering that *MPL1* expression, which is upregulated in response to GPA infestation, is strongest in the phloem, and *MPL1* expression from the phloem-specific *SUC2* promoter is sufficient to restore resistance to GPA in the *mpl1* background, it is likely that the effect of MPLI on limiting OPDA/JA accumulation is also exerted in the phloem.

**Fig. 7.**
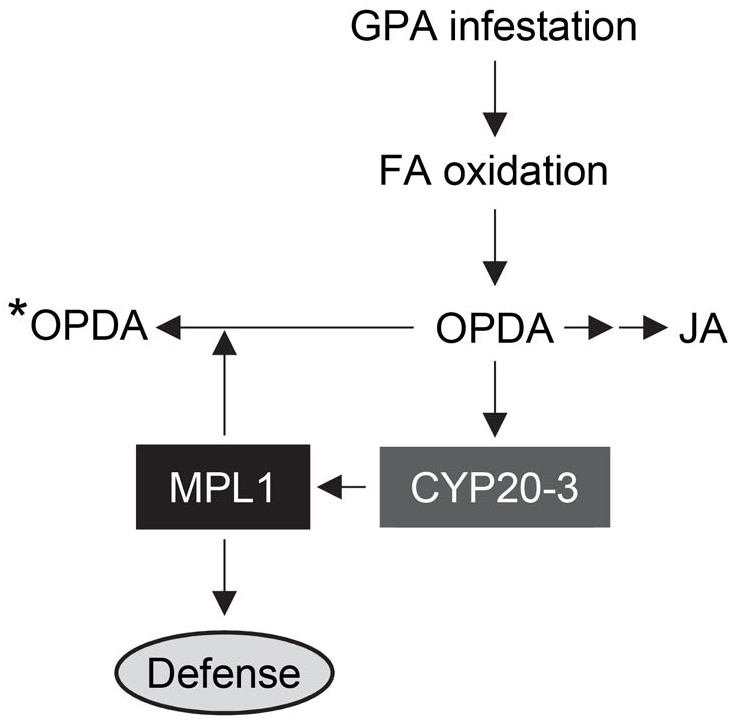
Model relating the interaction between *MPL1*, *CYP20-3* and OPDA in Arabidopsis defense against the green peach aphid (GPA). OPDA, which is synthesized in the plastids via enzymatic oxidation of fatty acids, is the precursor for jasmonic acid (JA) and its derivatives. OPDA also functions as a signaling metabolite. OPDA signaling requires *CYP20-3*. *CYP20-3* is shown to control expression of *MPL1,* which encodes a lipase that is required for limiting GPA infestation. *MPL1* in turn feedbacks to control OPDA level, presumably by limiting OPDA synthesis, or alternatively by promoting OPDA degradation or metabolism to other derivatives (indicated by *OPDA). Experiments with mutants defective in JA/JA-Ile biosynthesis from OPDA, and in JA/JA-Ile signaling, did not uncover significant contribution of JA-Ile signaling in defense against the GPA. In comparison, knockdown of *CYP20-3*, and genes encoding enzymes (13-LOX and AOS) involved in the synthesis of OPDA resulted in improved performance of GPA, supporting a role for OPDA signaling in defense against the GPA. The ability of OPDA to promote *MPL1* expression and resistance against the GPA, and of *MPL1* when constitutively expressed from the *35S* promoter in restoring resistance against the GPA in the *cyp20-3* mutant background supports this model.

Our analysis of GPA performance on *lox*(q), which lacks all for 13-LOX activities, and *aos*, which is deficient in allene oxide synthase, indicates that a product(s) of this pathway is required for Arabidopsis to limit GPA infestation. In contrast, GPA fecundity was not impacted on the *pxa1* and *acx1 acx3* mutants, which are defective the biosynthesis of JA (Theodoulou *et al*., 2005; Schilmiller *et al*., 2007), and *jar1*, which is defective in the synthesis of JA-Ile (Staswick and Tiryaki, 2004). In addition, GPA fecundity was comparable between the WT and *coi1*, which is defective in JA-Ile signaling (Katsir *et al*., 2008). These results suggest that while a 13-LOX and AOS-dependent product is required for limiting GPA fecundity, JA/JA-Ile signaling is not a major contributor to defense against the GPA. Similarly, Kettles *et al*. (2013) noted that GPA fecundity was not significantly different between the WT and *coi1* and *jar1*.

However, Ellis *et al*. (2002) and Mewis *et al*. (2006) reported that JA signaling contributes to Arabidopsis defense against the GPA. Ellis *et al*. (2002) observed that in comparison to the WT, GPA population size was smaller on the *cev1* mutant, which contains elevated JA content, and higher on the *coi1* and the *cev1 coi1* mutant plants. Similarly, Mewis *et al*. (2006) observed larger population of GPA on *coi1* compared to the WT plants. Ellis *et al*. (2002) further reported that resistance to the GPA was enhanced by JA application to the WT plant, but not the *coi1* mutant. We cannot rule out that this discrepancy between results on the involvement of JA in Arabidopsis-GPA interaction is due to differences in environmental conditions and/or potentially different GPA races/biotypes utilized in these studies; GPA is known to be a very variable species (van Emden *et al*., 1969). However, these studies (Ellis *et al*. 2002; Mewis *et al*., 2006; Kettles *et al*., 2013) did not set apart the individual contributions of JA and its precursor, OPDA, in providing resistance to aphids. Given that JA signaling feedbacks to upregulate *LOX2* and *AOS* expression and OPDA accumulation (Sasaki *et al*., 2001; Gleason *et al*., 2016), it is feasible that the impact of JA application on promoting resistance against GPA is associated with it promoting OPDA accumulation, as has been shown for Arabidopsis interaction with the root-knot nematode (*Meloidogyne hapla*) (Gleason *et al*., 2016). The MeJA application promoted resistance to root-knot nematode was mediated through OPDA and required *CYP20-3*, which is involved in OPDA signaling (Gleason *et al*., 2016). OPDA is a strong electrophile (Mueller and Berger, 2009) that is toxic to lepidopteran larvae (Vollenweider *et al*., 2000). However, we did not observe any direct toxic effect of OPDA on GPA when included in a synthetic diet. Instead, our observation that GPA fecundity was higher on *cyp20-3* compared to the WT, and OPDA application enhanced resistance to the GPA, suggests that OPDA signaling has a critical role in defense against the GPA. Similarly, OPDA promotes resistance against the GPA in radish seedlings (Guo *et al*., 2014). Also, in corn, OPDA has an important role in defense against the corn leaf aphid (Varsani *et al*., 2019). OPDA, but not JA, also promotes resistance against the brown planthopper in rice (Guo *et al*., 2014). Thus, the contribution of OPDA in defense against hemipterans that occupy a unique ecological niche via their ability to feed from the phloem, is not an uncommon phenomenon.

In conclusion, as summarized in Fig. 7, we provide evidence that an interplay between *MPL1*, OPDA, and *CYP20-3* influences Arabidopsis interaction with the GPA. *CYP20-3* controls expression of *MPL1,* which influences the accumulation in phloem sap of an antibiotic activity that inhibits aphid reproduction (Louis *et al*., 2010b). *MPL1* in turn feedbacks to control OPDA levels. Although the relationship between OPDA turnover and resistance to GPA is unclear, our results indicate that *CYP20-3* function, via its control of *MPL1* expression, is critical for controlling GPA infestation.

## Accession numbers

*ACX1 (At4g16760), ACX5* (At2g35690)*, AOS* (At5g42650)*, CML46* (At5g39670), *COI1* (At2g39940), *CYP20-3* (At3g62030), *EF-Tu, JAR1* (At2g46370), *LOX2* (At3g45140), *LOX3* (At1g17420), *LOX4* (At1g72520), *LOX6* (At1g67560), *MPL1* (At5g14180), *PXA1* (At4g39850), *SUC2* (At1g22710).

## Supplementary Data

Supplementary data are available at JXB online.

**Fig. S1.** Jasmonate biosynthesis pathway.

**Fig. S2**. *MPL1* contributes to Arabidopsis defense against the green peach aphid.

**Fig. S3.** *MPL1* and *CML46* expression in *lox*(q) plants.

**Table S1.** Primers used in this study.

## Acknowledgements

The authors would like to thank Drs. Sang Wook, Barbara Kunkel, and Ivo Feussner for the *cyp20-3*, *coi1-17*, and *lox2/3/4/6* quadruple mutants, respectively.

## Author Contribution

LA: conceptualized, designed and conducted experiments, analyzed data, and contributed to writing; SB: conducted experiments and analyzed data; HAM: conceptualized, designed and conducted experiments and analyzed data; MT: conceptualized, designed and conducted experiments and analyzed data; JL: conceptualized, designed and conducted experiments, analyzed data, and contributed to writing; VJM: conceptualized, designed and conducted experiments and analyzed data; JK: designed and conducted experiments and analyzed data; ZC: conducted experiments and analyzed data; JS: conceptualized, administered and supervised the research, contributed to funding acquisition and writing.

## Data Availability

All data supporting the findings of this study are available within the paper and within its supplementary materials published online.

