## Supplemental Figures S1 - S3 for "Interplay between *MYZUS PERSICAE-INDUCED LIPASE 1* and OPDA signaling in controlling green peach aphid infestation on *Arabidopsis thaliana*"

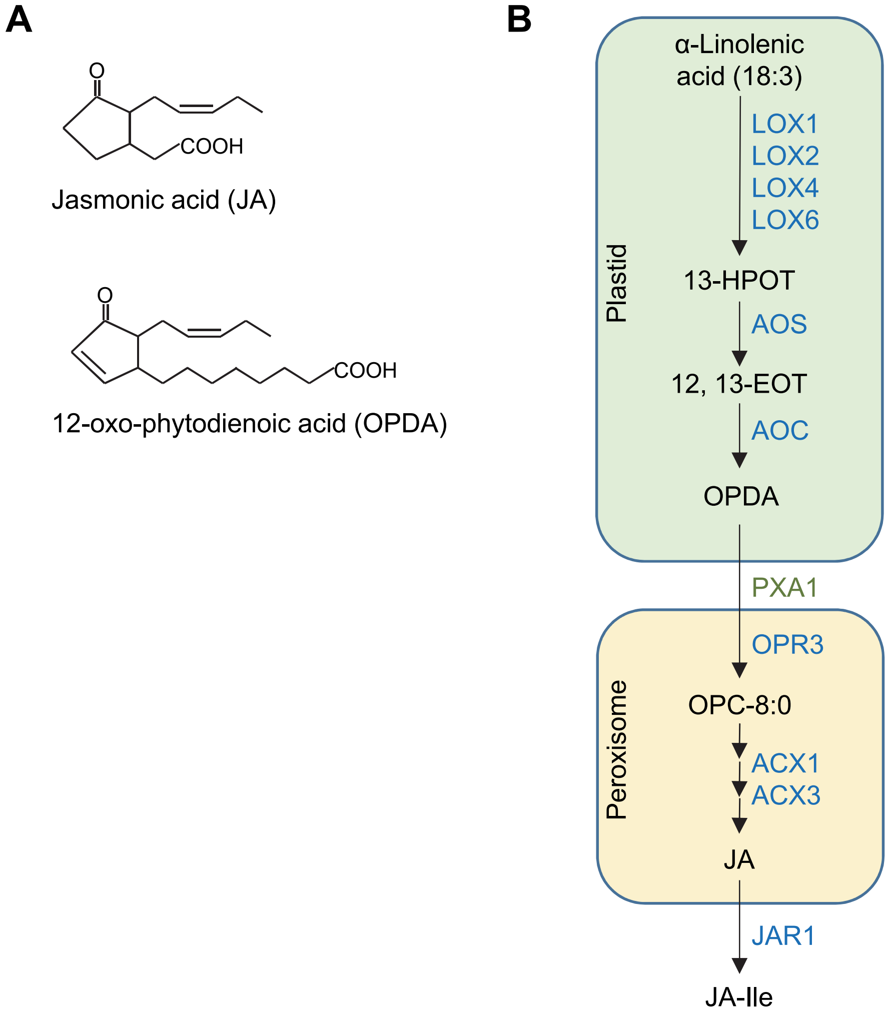


**Fig. S1.** Jasmonate biosynthesis pathway. **(A)** Jasmonic acid (JA) and 12-oxo-phytodienoic acid (OPDA). **(B)** Biosynthesis pathway for OPDA, JA and JA-isoleucine (JA-Ile). The biosynthesis of jasmonates involves multiple steps with the dioxygenation of the polyunsaturated fatty acid α-linolenic acid (C18:3), catalyzed by 13-lipoxygenase (13-LOX) to yield the corresponding 13-hydroperoxytrienoic acid (13-HPOT). 13-HPOT is sequentially acted upon by allene oxide synthase (AOS) and allene oxide cyclase to yield 12-oxo-phytodienoic acid (OPDA). The steps leading to the synthesis of OPDA occur in the plastid (green box). OPDA is subsequently transported from the plastids to the peroxisome (yellow box), a step that involves the peroxisomal ABC transporter1 (PXA1). In the peroxisome, the reduction of OPDA by OPDA reductase 3 (OPR3) yields 3-oxo-2 (2′(Z)-pentenyl)-cyclopentane-1-octanoic acid (OPC-8:0), which is subsequently channeled through three rounds of *β*-oxidation that involves the acyl-CoA oxidases ACX1 and ACX3 to yield jasmonic acid (JA). JA is then transported to the cytosol where it is derivatized to JA-Ile by the jasmonoyl-isoleucine synthetase, JAR1.

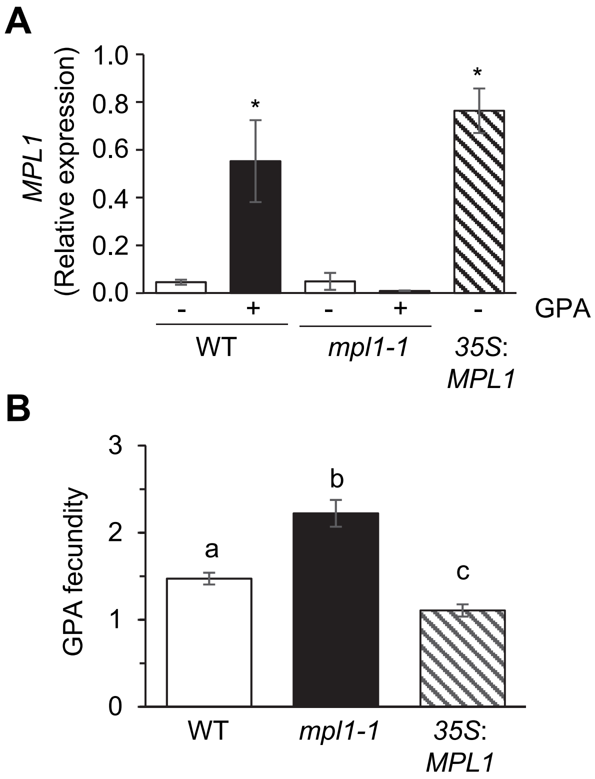


**Fig. S2**. *MPL1* contributes to Arabidopsis defense against the green peach aphid. **(A)** Upregulation of *MPL1* expression in response to GPA infestation. *MPL1* transcript levels (x $10^{-2}$) relative to the elongation factor gene At1g07920 at 24 hpi with GPA (+GPA) on leaves of wild-type (WT) Arabidopsis accession Columbia and the *mpl1-1* mutant plants. Leaves harvested at the same time from uninfested (-GPA) WT, *mpl1-1* and *35S:MPL1* plants provided the controls. The *35S:MPL1* plants constitutively express *MPL1* from the *Cauliflower mosaic virus 35S* promoter. Error bars represent $\pm$ SE (n=3). Asterisks indicate values that are significantly higher than the uninfested WT (P<0.05; *t*-test). (**B**) GPA fecundity on the WT, *mpl1-1* and *35S:MPL1* plants. Error bars represent $\pm$ SE (n=15). Different letters above bars represent values that are significantly different from each other (P<0.05; ANOVA).

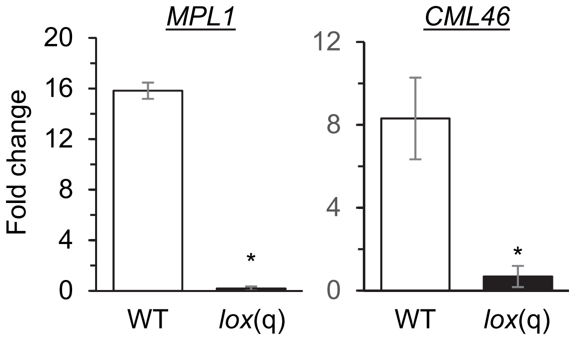


**Fig. S3.** *MPL1* and *CML46* expression in *lox*(q) plants. Fold change in expression of *MPL1* and *CML46* in the GPA-infested relative to the uninfested WT and *lox*(q) plants. Expression was monitored at 24 hpi. Error bars represent ± SE (n=3). Asterisks indicate values that are significantly different from the WT (P<0.05; *t*-test).
