## Supplemental Table S1 for "Interplay between *MYZUS PERSICAE-INDUCED LIPASE 1* and OPDA signaling in controlling green peach aphid infestation on *Arabidopsis thaliana*"

**Table S1.** Primers used in this study

| Primer Name | Gene | Primer Sequence (5'→3') | Purpose |
|---|---|---|---|
| CYP20-3-F1 | At3g62030 | GGGTTCGGTGTTAAGTCGCAATTAG | Genotyping |
| CYP20-3-R1 | At3g62030 | CTAACCATGAAGTCTGCAC | Genotyping |
| MPL1-F1 | At5g14180 | CGGGGATCGACTGTTATGAT | qPCR |
| MPL1-R1 | At5g14180 | TGCGATCACTGCTTCCATAG | qPCR |
| MPL1-CDS-Stop-R1 | At5g14180 | TCAAGCTTGTCGTTTGAAAAAGG | Genotyping |
| EF-Tu-F1 | At1g07920 | TGCCGCAGGTGAATCAAAGG | qPCR |
| EF-Tu-R1 | At1g07920 | CCCAATTACGAGAACAACGCTCTG | qPCR |
| CML46-F1 | At5g39670 | CCTCTTGTTCAACACCAGCA | qPCR |
| CML46-R1 | At5g39670 | GCTTTAAGCCAAGGATTGTCA | qPCR |
| SUC2p-F1 | At1g22710 | TCTTCCTCTTCCTCCACCA | Genotyping |
| 35S-F1 | | GTGATATCTCCACTGACGTAAGG | Genotyping |
| T-DNA-LB | | GGTTCACGTAGTGGGCCATC | Genotyping |
